# Spatial self-organization of confined bacterial suspensions

**DOI:** 10.1101/2025.02.20.639202

**Authors:** Babak Vajdi Hokmabad, Alejandro Martínez-Calvo, Sebastian Gonzalez La Corte, Sujit S. Datta

## Abstract

Lab studies of bacteria usually focus on cells in spatially-extended, nutrient-replete settings, such as in liquid cultures and on agar surfaces. By contrast, many biological and environmental settings—ranging from mucus in the body to ocean sediments and the soil beneath our feet—feature multicellular bacterial populations that are confined to tight spots where essential metabolic substrates (e.g., oxygen) are scarce. What influence does such confinement have on a bacterial population? Here, we address this question by studying suspensions of motile *Escherichia coli* confined to quasi two-dimensional (2D) droplets. We find that when the droplet size and cell concentration are both large enough, the initially-uniform suspension spatially self-organizes into a concentrated, immotile inner “core” that coexists with a more dilute, highly-motile surrounding “shell”. By simultaneously measuring cell concentration, oxygen concentration, and motility-generated fluid flow, we show that this behavior arises from the interplay between oxygen transport through the droplet from its boundary, uptake by the cells, and corresponding changes in their motility in response to oxygen variations. Furthermore, we use theory and simulations to develop quantitative principles describing this interplay—establishing a biophysical framework that unifies all our experimental observations. Our work thereby sheds new light on the rich collective behaviors that emerge for bacterial populations, and other forms of chemically-reactive living and active matter, in confined environments, and provides a way to predict and control these behaviors more broadly.

---

Bacteria often inhabit confined spaces, ranging from the thin mucus layer that coats the internal organs of most animals [1–7] to the pores in subsurface soils and sediments [8– 14]. In such environments, multicellular consumption of scarce metabolites can create strong gradients across a bacterial population [15–22]. These metabolite gradients, in turn, can give rise to physiological heterogeneity throughout the population—even if its constituents are all genetically identical—since distinct cells experience different chemical microenvironments and thus behave differently [21–25].

This form of “phenotypic differentiation” can have profound implications for the organization and functioning of a population. For example, oxygen (O_2_) consumption by aerobes can establish anoxic niches for obligate anaerobes, enabling both cell types to stably coexist [26–30]—facilitating the ability of such mixed populations to cycle nutrients in soil [28], perform key metabolic processes in the gut [31], and remove pollutants from dirty water [32]. As a result, the influence of collectively-generated metabolite gradients on confined bacterial populations is well-studied in the context of variations in gene expression, metabolic activity, chemical signaling, and growth/proliferation [11, 16, 33–42].

Less studied, however, is how such gradients shape, and are shaped by, cellular motility—another fundamental characteristic of many bacteria. Studies of *unconfined* populations of motile cells have revealed that they migrate as coherent fronts in response to self-generated gradients of chemoattractants/O_2_, a phenomenon known as collective chemotaxis/aerotaxis [43–57]. And remarkably, cells of different motility characteristics self-sort within such a front in response to the shape of the gradient [44]. The implications of self-generated metabolite gradients on *confined* populations of motile bacteria are, however, less explored.

Here, we report a distinct form of phenotypic differentiation that arises in confined populations of motile aerobic bacteria. We study initially-uniform suspensions of swimming *E. coli* confined to quasi-2D droplets. When the droplet is sufficiently large and concentrated, we find that the population spontaneously self-organizes into a concentrated inner core of immotile cells surrounded by a more dilute annular shell of highly-motile cells—even though all the cells are genetically identical. By integrating experiments, theory, and simulations, we show that this phenomenon is governed by the interplay between cellular aerobic respiration, O_2_ transport and availability, and cellular motility. In particular, O_2_ supplied from the droplet boundary is taken up by the swimming cells, eventually becoming depleted and causing cells in the core to rapidly lose motility—as has been studied in other contexts [29, 58–60]. In some cases, this spatial self-organization is permanent. In others, the population continues to restructure itself over longer time scales: aerotaxis enables the swimming cells to reshape the O_2_ gradient in turn and promote O_2_ influx, causing the core to eventually disappear. We establish quantitative principles that describe the conditions under which anoxic core formation arises and whether it is transient or permanent—in excellent agreement with all our experiments and simulations. Hence, this work sheds new light on the fascinating behaviors that can emerge from the interplay of confinement, collectively-generated metabolite gradients, and cellular motility. In doing so, it provides principles for predicting and controlling the spatial organization of bacterial populations, and other forms of chemically-reactive living and active matter, more broadly.

## RESULTS

### Confined *E. coli* suspensions spontaneously self-organize in three distinct ways

To explore the collective dynamics of confined motile bacteria, we sandwich droplets of suspensions of swimming *E. coli* between two flat glass plates (Fig. **1**a) in a humid environment to minimize evaporation. The gap between the plates is 130 µm high, much smaller than the droplet radius *R*, which we set to be between 0.3 and 5.5 mm. The plates themselves are O_2_-impermeable, and so, O_2_ can only enter from the air-liquid interface at the droplet perimeter. To avoid confounding effects, we use a buffer that only contains non-metabolized salts; hence, the cells do not proliferate, but instead, use their internal resources to maintain motility over many hours and only respond to variations in O_2_ (Fig. S1, [59]). The cells are initially uniformly dispersed at a concentration *c*_cell,0_. They constitutively express green fluorescent protein, enabling us to map subsequent variations in the depth (*z*)-integrated cellular concentration *c*_cell_(**r**, *t*) throughout each droplet via confocal microscopy using both bright-field and fluorescence imaging; **r** and *t* represent the radial coordinate and time, respectively.

**Fig. 1.**
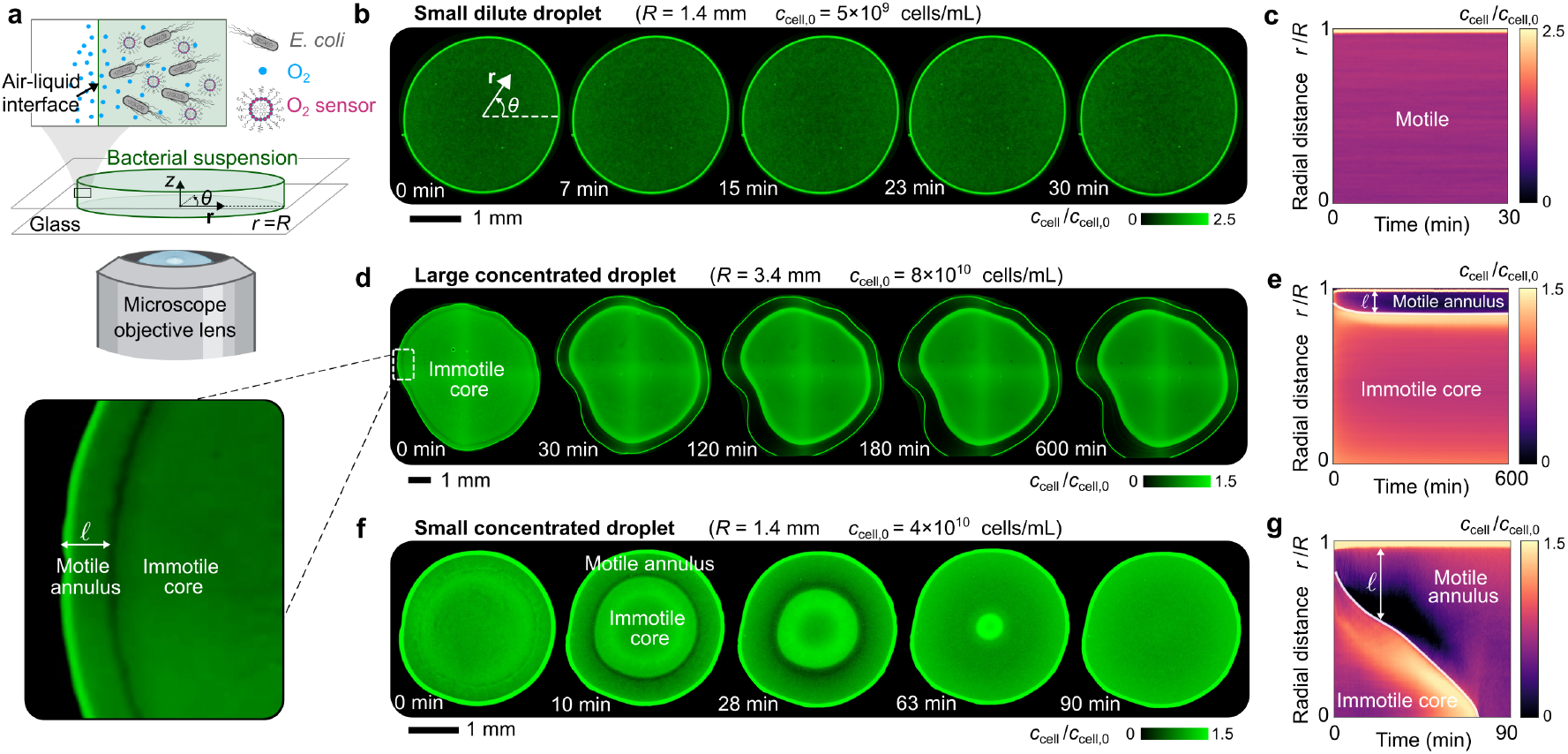
Spontaneous spatial self-organization of confined bacterial suspensions. **a**, Schematic of the experimental setup. A droplet containing swimming fluorescent *E. coli* (grey) is sandwiched between two flat glass plates separated by a gas-permeable spacer and maintained in a humid environment at 30°C. Oxygen (blue) diffuses in from the air-liquid interface, and is detected using a dissolved fluorescent probe encapsulated in phospholipid micelles (pink). **b, d, f**, Depth-averaged bright-field micrographs showing the time evolution of bacterial suspensions (Movies S1-S4) for (**b**-**c**) a small dilute droplet, (**d**-**e**) a large concentrated droplet, and (**f**-**g**) a small concentrated droplet, corresponding to 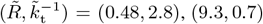, and (2.7, 1), respectively. The kymographs in **c, e, g**, represent these dynamics by showing the azimuthally-averaged cell concentration *c*_cell_, normalized by its initial spatially-uniform value *c*_cell,0_, as a function of radial position *r* and time *t*. The small dilute droplet remains nearly uniform; by contrast, in more concentrated droplets a concentrated core of immotile cells forms, surrounded by a less-concentrated annular shell of motile cells. In large droplets, this core forms permanently, while in small droplets, this core shrinks and eventually disappears.

We start with a small (*R* = 1.4 mm) and dilute (*c*_cell,0_ = 5 × 10^9^ cells/mL) droplet. In this case, the entire population remains motile for the duration of the experiment, as shown in Movie S1. Almost all the cells are uniformly dispersed, with a minute fraction localized at the air-liquid interface, as shown by the bright thin band along the droplet perimeter in Figs. **1**b-c.

We observe strikingly different behavior in the case of a larger (*R* = 3.4 mm) and more concentrated (*c*_cell,0_ = 8 × 10^10^ cells/mL) droplet. Instead of being uniformly dispersed, the cells rapidly self-organize into two distinct domains (Movie S2): a large, concentrated inner core surrounded by a thinner, more dilute annular shell, as shown by the brighter and darker regions, respectively, in Fig. **1**d. A minute fraction of cells again localizes at the air-liquid interface. The cells in the different domains have markedly different motility characteristics; in the core, cells are completely immotile, whereas in the annulus, they remain fully motile, as shown in Movie S3. Interestingly, the core shrinks slightly, and cells accumulate near its periphery and are depleted from the surrounding annulus, during the first ∼100 min. The core then reaches a steady-state size, as shown in Fig. **1**e. Correspondingly, the surrounding annulus has a nearly constant width *𝓁* ∼300 µm, following the shape of the droplet boundary and even tracking slight imperfections in droplet shape (Fig. **1**d).

We observe yet another different behavior in the case of a small (*R* = 1.4 mm) but still concentrated (*c*_cell,0_ = 4 × 10^10^ cells/mL) droplet. The cells again rapidly self-organize into a concentrated inner core, in which they are immotile, surrounded by a more dilute annular shell in which they remain motile (Movie S4). Moreover, cells in the core again appear to accumulate near its periphery and are depleted from the surrounding annulus. However, we observe two key differences from the previous case. First, a larger fraction of the motile cells localizes at the air-liquid interface, as shown by the thicker bright band along the droplet perimeter in Fig. **1**f. Second, the self-organization into a core-shell structure is transient. As time progresses, the core appears to be “eroded” by the shell, progressively shrinking until it ultimately disappears after ∼70 min (Figs. **1**f-g), reaching a final steady state similar to the first case shown in Figs. **1**b-c.

### Bacterial self-organization is governed by the interplay between oxygen transport and availability, cellular respiration, and motility

Why do confined, concentrated bacterial populations self-organize into a core-shell structure? And why is this spatial organization permanent in some cases and transient in others? Close inspection of the cellular swimming patterns in concentrated droplets (Movie S3), obtained using high-magnification imaging 30 µm above the bottom glass plate, provides a clue. Just outside the core, the time-averaged velocity field associated with bacterial swimming splits into two coherent, radially directed streams (Figs. S2-S3). The stream closer to the core is directed radially inward (pink in Fig. S2), indicating motile cells that swim inward and rapidly lose motility, causing them to accumulate at the core periphery—that is, the core acts as a *sink* for cells. The outer stream is directed radially outward (green in Fig. S2), resulting in eventual depletion of cells from the region surrounding the core and localization of cells at the air-liquid interface instead. These rapid changes in motility suggest that a metabolic shift is at play — presumably in response to spatial variations in dissolved O_2_, given that it is the only exogenous metabolite available to the cells in our experiments, and the cells use it to perform aerobic respiration. Indeed, prior studies [58, 59] have shown that *E. coli* rapidly loses motility, just as we observe in the core, when deprived of O_2_. Other studies have shown that when O_2_ is available, *E. coli* biases its swimming toward regions of higher O_2_ availability [45], just as we observe near the air-liquid interface. Moreover, large, concentrated droplets that are not confined by an overlying glass plate as in our experiments, but are instead open to atmospheric O_2_, do not exhibit core formation [61]. Taken altogether, these observations suggest that the self-organization of confined *E. coli* suspensions is governed by the interplay between oxygen transport and availability, cellular respiration, and motility.

To test this idea, we use a fluorescent probe [58] to directly map variations in dissolved O_2_ throughout a confined droplet (Movie S5). The droplet is small and concentrated, and so, exhibits transient core-shell formation as in Fig. **1**f. Immediately after the start of the experiment, we observe a radial gradient in O_2_ concentration, 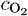, progressively decreasing from its saturation value 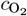,sat at the outer air-liquid interface inward and eventually reaching zero at the periphery of the core (Fig. **2**a). That is, the formation of a core of immotile cells is indeed coincident with the establishment of an anoxic region inside the droplet—which arises presumably because of uptake by the swimming cells in the surrounding annular shell. If this conjecture is correct, then the annulus width *𝓁* should be set by the distance over which O_2_ supplied from the droplet boundary can diffuse radially inward before being fully consumed by the swimming cells: 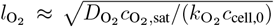 [62], where 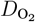 is the diffusivity of O_2_ and 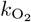 is the maximal O_2_ uptake rate per cell. We directly test this prediction by varying *c*_cell,0_ over an order of magnitude and measuring the corresponding *𝓁*. As shown by the red points in Fig. **2**b, our experimental measurements confirm this prediction. As an additional test, we perform the same experiments, but for cells dispersed in a nutrient-rich liquid—in which active metabolism of nutrients increases the rate of O_2_ uptake, 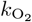. Our measurements again confirm this prediction, as indicated by the blue points in Fig. **2**b. These data quantitatively establish that, in large and concentrated droplets, O_2_ depletion due to uptake by swimming cells in the outer annular shell causes inner cells to lose motility, sediment, and form a concentrated core.

**Fig. 2.**
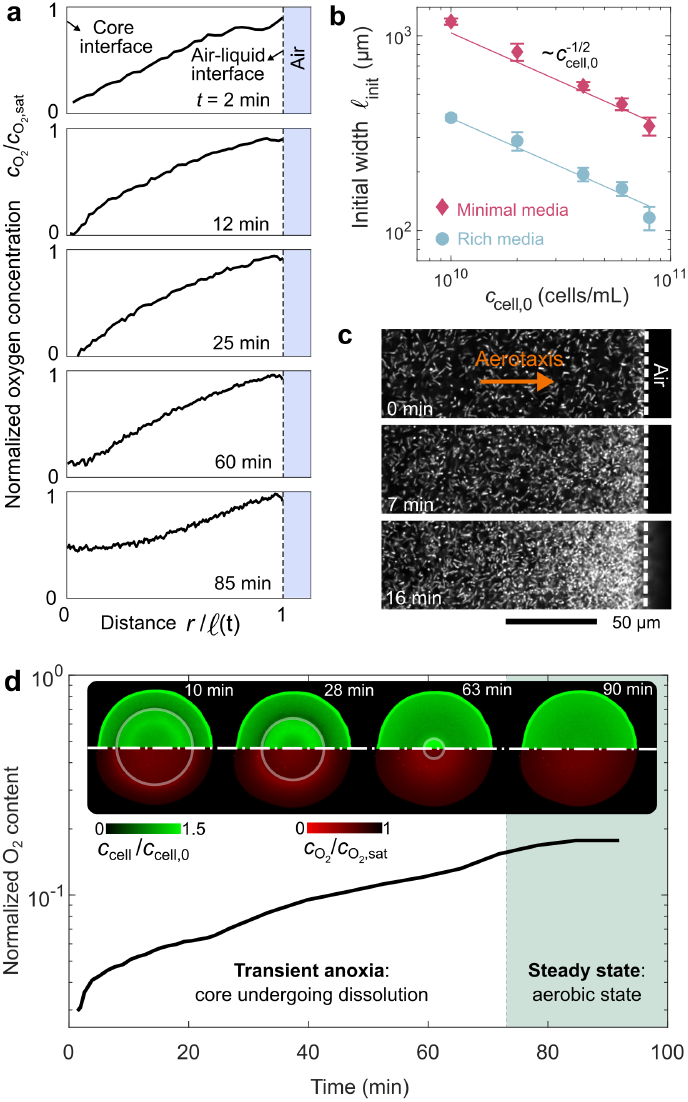
Bacteria regulate oxygen transport through uptake and by accumulating at the air-liquid interface. **a**, Evolution of the azimuthally-averaged profile of oxygen concentration 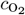, normalized by its saturation concentration 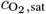, as a function of radial position across the annular shell, *r*; *𝓁* is the time (*t*)-dependent annulus width. This measurement is for a droplet with *R* = 1.4 mm and *c*_cell,0_ = 4 × 10^10^cells*/*mL, corresponding to 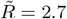 and 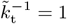. Cellular uptake creates an O_2_ gradient, with 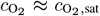 at the air-liquid interface and 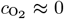 at the periphery of the core. Over time, however, influx of O_2_ causes it to re-aerate the core. **b**, Variation of the initial width of the motile annulus *𝓁*_init_ after core formation with increasing cell concentration *c*_cell,0_ in minimal media (BMB, red) and in nutrient-rich media (LB Broth, blue). The lines show excellent agreement with our theoretical prediction for the oxygen penetration length 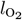, with uptake rates 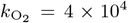 molecules s^*−*1^ cell^*−*1^ in minimal media and 3 × 10^5^ molecules s^*−*1^cell^*−*1^ in nutrient-rich media, consistent with prior measurements [44, 58, 62]. **c**, High magnification fluorescent confocal micrographs taken at a planar optical section in the middle of the channel (Movie S6) reveal cell accumulation near the air-liquid interface with time due to aerotaxis. Experiment was done at *R* = 1.4 mm and *c*_cell,0_ = 10^10^cells*/*mL, corresponding to 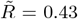 and 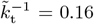. **d**, Time evolution of the total O_2_ content of the droplet in **a** obtained by integrating the measured O_2_ concentration over the annulus, normalized by the total O_2_ content at the saturation condition. Inset shows micrographs of the profiles of cells (top half) and dissolved O_2_ (bottom half) throughout the droplet (Movie S5). The bacteria are imaged via depth-averaged bright-field microscopy and the dissolved O_2_ are imaged using fluorescent confocal microscopy from an optical slice that is ∼ 00µm thick. Note that the dimming of the O_2_ signal within the anoxic core area is due to light scattering from the concentrated layer of bacteria sedimented on the bottom surface. Our measurements show that after the core forms, enhanced O_2_ influx is concomitant with core shrinkage (“transient anoxia”), ultimately causing it to disappear and marking the final aerobic steady state.

Having identified the mechanism of core formation, we next ask: Why, in some cases, is the core eventually eroded away? Examining the O_2_ and cell concentration profiles simultaneously, which shows how they are coupled, helps to address this question. Not only does O_2_ uptake cause the core of the droplet to become anoxic, but it also establishes a radial gradient through the annular shell (Fig. **2**a) that the swimming cells respond to via aerotaxis. An example is shown in Fig. **2**c (Movie S6). As these swimming cells accumulate near the air-liquid interface, their respiration continues to reshape this O_2_ gradient, promoting its further influx from the droplet periphery. Our 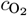 measurements directly confirm this picture. We integrate them both radially (*r*) and azimuthally (*θ*) to determine the total amount of O_2_ in the droplet: 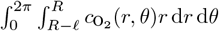. As shown in Fig. **2**d, O_2_ content monotonically increases over time, concurrent with the erosion of the anoxic core. This process persists until the core disappears altogether (inset, Fig. **2**c), resulting in a final aerobic steady state. By biasing its distribution within the outer annulus via aerotaxis, the motile subpopulation of cells promotes O_2_ influx across the air-liquid interface; as a result, core formation is only transient.

### Quantitative principles underlying bacterial self-organization

To further rationalize the experimental observations, we construct a continuum model that describes the interplay between oxygen transport and availability, cellular respiration, and motility in confined bacterial suspensions. Given that the experimental droplets are quasi-2D, for simplicity, we consider depth (*z*)-integrated quantities in an axisymmetric 2D system. The model is summarized in Fig. **3**a. Eq. (1a) relates local changes in O_2_ levels to its diffusion and uptake by the motile cells, which in turn is modulated by oxygen availability relative to the characteristic concentration *K* = 1 µM via Michaelis-Menten kinetics. Eq. (1b) provides a boundary condition relating O_2_ influx at the air-liquid interface, 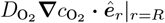, to the difference between 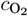 and 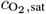 through the mass transfer coefficient *k*_t_; here, ***ê***_*r*_ is the unit radial vector. Finally, Eq. (1c) relates local changes in bacterial concentration to their motility, which has two components— undirected random motion and biased random motion up an O_2_ gradient. The former can be described as a diffusive process with an “active” diffusivity *D*_cell_. The latter is described using the monotonically increasing function 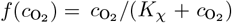, which describes the ability of the cells to sense variations in O_2_ based on first-order kinetics of receptor binding with a dissociation constant *K*_*χ*_ = 7 µM [46], along with the aerotactic coefficient *χ* describing their ability to move up the sensed gradient. Therefore, 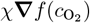 can be thought of as an aerotactic velocity, and when multiplied by *c*_cell_ describes the aerotactic bacterial flux. Importantly, to account for the loss of cellular motility when deprived of O_2_, both motility parameters *D*_cell_ and *χ* become negligible when 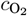 drops below a critical value *c*_crit_ ≪ *K*_*χ*_ [58].

**Fig. 3.**
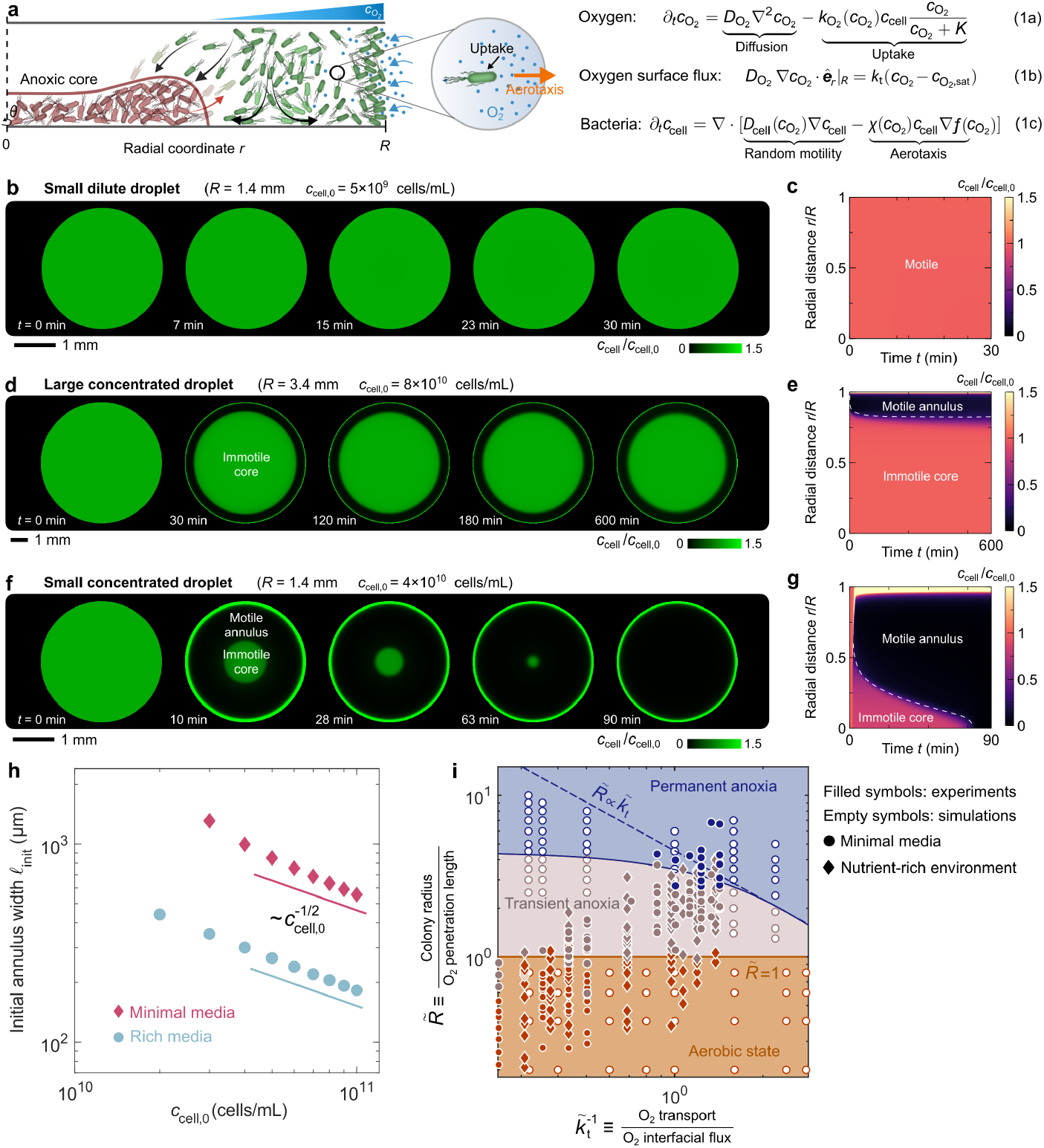
Theoretical model recapitulates the experimental observations and unifies them in a morphological state diagram. **a**, Schematic of the biophysical principles underlying core formation and shrinkage. Motile bacteria in the annulus (green) swim either randomly into the anoxic core (left-pointing arrows), where they lose motility (red), settle, and concentrate themselves, or they move up the O_2_ gradient via aerotaxis (right-pointing arrow) and accumulate at the air-liquid interface. They do not block the influx of O_2_, which is much smaller [63] than the interstices between cells. Conversely, their uptake promotes further O_2_ influx, which can cause cells at the core periphery to become motile again (right-pointing red arrow), driving core erosion and shrinkage. These feedbacks between O_2_ transport and influx, its uptake by cells, and cellular motility are quantified by Equations (1a-c), along with the condition that at the air-liquid interface, there is no flux of bacteria from the liquid phase into the gas i.e., 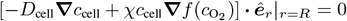. To describe the loss of bacterial motility at low O_2_ concentration, we use the phenomenological expressions 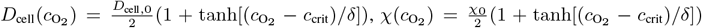, and 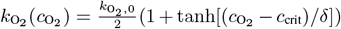, where *δ* controls the sharpness of the transition at 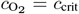. **b-g**, Numerical simulations of the time evolution of bacterial suspensions (Movies S7-S9) for the exact same conditions as the experiments in Fig. **1**b-g. As in the experiments, the small dilute droplet is uniform, while more concentrated droplets have an inner concentrated core of immotile cells surrounded by a less-concentrated annular shell of motile cells that is either permanent or transient. **h**, The simulations also recapitulate the experimental measurements of the initial annulus width shown in Fig. **1**h. **i**, State diagram is spanned by the two dimensionless parameters 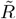 and 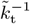, where the former compares the droplet radius to the O_2_ penetration length, and the latter compares the rates of O_2_ transport via diffusion and influx from the air-liquid interface. Filled and empty symbols show experimental and simulation results, respectively. Circles and diamonds indicate minimal media and nutrient-rich conditions, respectively. The solid dark orange line shows our theoretical prediction for the boundary between the aerobic state and transient anoxia, 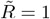. The dashed dark blue line shows our theoretical prediction for the boundary between the transient and permanent states of anoxia, 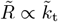, which assumes that O_2_ is not saturated at the air-liquid interface, appropriate for 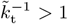; the solid dark blue line shows this boundary without making this assumption, as determined by simulations. In all cases, we find excellent agreement between theory, simulations, and experiments.

Remarkably, numerical simulations of this minimal model show that it recapitulates all the key features of the experimental observations. When the droplet is small and dilute, it remains fully aerobic; therefore, the entire population remains motile and nearly uniformly dispersed, as shown in Fig. **3**b-c and Movie S7 (compare to Fig. **1**b-c and Movie S1). When the droplet is larger and more concentrated, the cells self-organize into a concentrated immotile core—which remains permanently anoxic—surrounded by a dilute motile annular shell of width *𝓁* ∼ 300 µm, as shown in Fig. **3**d-e and Movie S8 (compare to Fig. **1**d-e and Movie S2). Moreover, we find the same dependence of this annular shell width on *c*_cell,0_ as in the experiments, as shown in Fig. **3**h (compare to Fig. **2**b). Finally, when the droplet is small but concentrated, a larger fraction of motile cells localizes at the air-liquid interface, and anoxic core formation is transient, just as in the experiments, as shown in Fig. **3**f-g and Movie S9 (compare to Fig. **1**f-g and Movie S4). These results indicate that the biophysical picture described in Fig. **3**a captures the essential processes underlying the spatial self-organization of confined bacterial suspensions.

Analysis of the model also enables us to establish quantitative principles describing the conditions under which each of the three states observed—aerobic, permanently anoxic core-shell, and transiently anoxic core-shell—arises. To do so, we first non-dimensionalize the governing equations in Fig. **3**a, revealing two dimensionless parameters that describe a confined bacterial suspension (*Materials and Methods*): 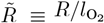, which compares the size of the droplet to the distance O_2_ penetrates into it before being fully consumed, and 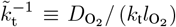, which compares the rates of O_2_ transport via diffusion, 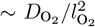, and influx from the air-liquid interface, 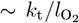. The conditions demarcating transitions between the three states of a confined bacterial suspension can then be expressed in terms of these two parameters. Indeed, a necessary condition for an anoxic core to form, whether transiently or permanently, is that 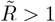; otherwise, O_2_ can freely penetrate throughout the droplet, and the aerobic, core-free state is maintained. Furthermore, when 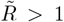, we expect that anoxic core formation is permanent when O_2_ uptake by the entire population, 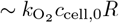, outpaces influx from the air-liquid interface, which we estimate as 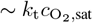 when O_2_ influx is not limited by saturation at the air-liquid interface. That is, we expect that for 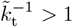, anoxic core formation is permanent when 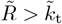.

To test these predictions quantitatively, we perform 256 different experiments and 97 different simulations, systematically exploring a broad range of droplet radii *R* and cell concentrations *c*_cell,0_ in both minimal and nutrient-rich liquid media. The results are summarized by the points in Fig. **3**i, which represents a morphological state diagram spanned by the two dimensionless control parameters revealed by our analysis, 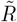 and 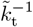. The three different colors represent the three different states observed: dark orange represents conditions under which the droplet remains in the aerobic, fully-motile state, while grey-orange and blue represent conditions under which the droplet self-organizes into an anoxic, immotile core surrounded by an aerobic, motile shell either transiently or permanently, respectively. Points in these three different states cluster in different regions of the diagram, confirming the utility of 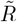 and 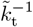 as dimensionless control parameters. The transition from the aerobic state to that of anoxic core formation is described well by the predicted 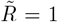, shown by the horizontal dark orange line. Moreover, when 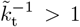, the transition from transient to permanent anoxic core formation is also described well by the predicted 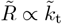 with a prefactor ≈4, shown by the dashed blue line. Conversely, for 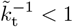, O_2_ reaches its saturation concentration at the air-liquid interface (*Materials and Methods*), constraining its influx and removing the dependence of the transition from transient to permanent anoxic core formation on 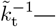as expected, and as quantitatively shown by the solid blue line. Thus, our integrated experiments, theoretical analysis, and simulations establish biophysical principles that quantitatively describe how confined bacterial suspensions self-organize across different levels of confinement.

### Viscoelasticity generates strong fluctuations during core erosion

In many cases, bacteria are confined to polymeric fluids, such as mucus in the body [64–67], exopolymers in the ocean, and extracellular polymeric matrices in biofilms [68]. Recent work has shown that the viscoelasticity of these fluids can alter how individual motile cells swim in unusual ways—for example, by markedly increasing the swimming speed [69–74] and enhancing fluid-mediated interactions between cells [61, 75]. Thus, as a final exploration of bacterial self-organization in confinement, we ask: How do interactions with extracellular polymers influence this process?

To address this question, we repeat the experiment shown in Fig. **1**f, but with 5 MDa polyethylene oxide (PEO) added to the fluid at a semidilute concentration (0.2 wt.%) comparable to that of mucus in the body. This synthetic polymer is chemically inert, uncharged, nonadsorbing, and similar in size to many biological mucins. Our measurements confirm that cell swimming is enhanced in this polymeric fluid, with a ∼ 4-fold increase in *D*_cell_ compared to the polymerfree case (Table S1). The experiment is characterized by 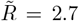 and 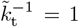, and so, we expect that just like the polymer-free case, an anoxic immotile core forms, surrounded by an oxygenated motile annular shell, and then disappears (Fig. **3**i). However, simulation indicates that because of their enhanced motility, cells accumulate at the air-liquid interface more rapidly than in the polymer-free case, promoting O_2_ influx and causing the core to shrink faster (compare Movies S4 and S10).

Our experimental observations confirm these predictions, but also reveal additional rich dynamics imparted by fluid viscoelasticity. As shown in Movie S11 and Fig. **4**, a concentrated immotile core initially forms (*t* ≲ 300 s), with cells accumulating near its periphery and depleting from the surrounding annulus. However, unlike the polymer-free case, we observe large (∼100 µm) coherent swirls generated by the enhanced cellular swimming [60, 76] at the periphery of the core. As time progresses (*t* ≳ 300 s), these swirls propagate into and throughout the core. Concomitantly, a large fraction of motile cells—much larger than in the polymer-free case, as seen by comparing the third panels of Figs. **4** and **1**f—localizes at the air-liquid interface. As before, the increased O_2_ influx then causes the core to continue to shrink until it has eventually disappeared (*t* ≳ 2900 s)—faster than in the polymer-free case of Fig. **1**f. Additionally, unlike the polymer-free case, the cells are not uniformly distributed throughout the droplet at the end of the experiment; rather, they are depleted from the interior and instead remain localized at the air-liquid interface, as shown by the thick band in the last panel of Fig. **4**. Taken altogether, these observations demonstrate that our central finding—that confined bacterial suspensions self-organize into core-shell structures due to O_2_ limitation—is more general, manifesting in complex fluids akin to those encountered in many biological settings. Further unraveling the additional dynamics imparted by fluid viscoelasticity will be a fascinating direction for future work.

**Fig. 4.**
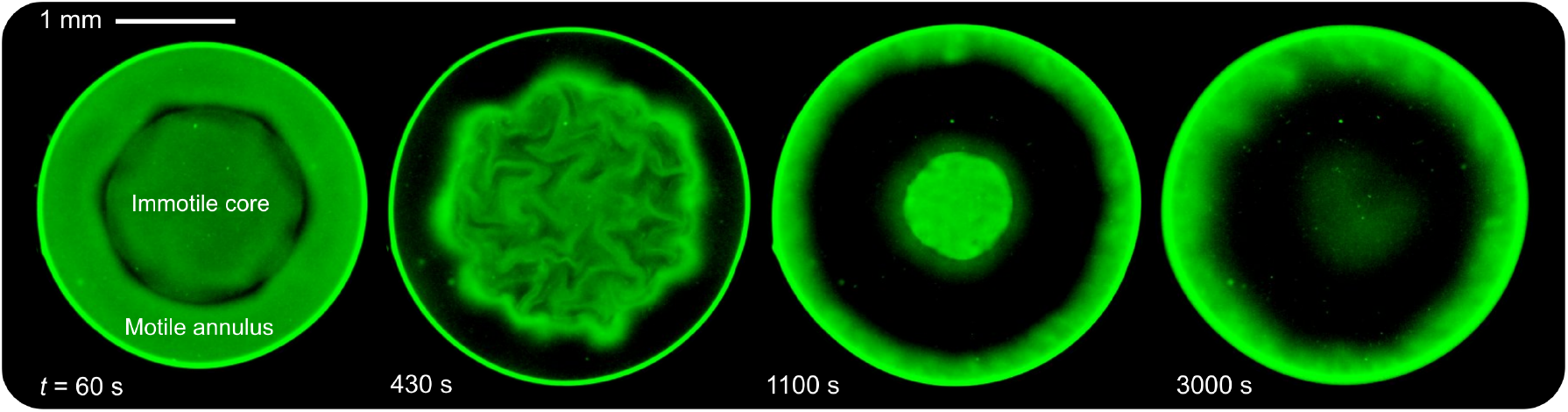
Fluid viscoelasticity imparts strong and coherent fluctuations during bacterial self-organization. Depth-averaged bright-field microscopy micrographs showing the time evolution of a bacterial suspension (Movie S11) with *R* = 1.4 mm and *c*_cell,0_ = 4 × 10^10^cells*/*mL, corresponding to 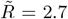 and 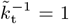, in a viscoelastic solution of 5 MDa PEO at 0.2wt.% (≈ 4 times its overlap concentration). As our theory predicts, an immotile core forms, surrounded by a motile annulus. However, the swimming cells in the annulus generate coherent swirls that mix the core. More cells also localize at the air-liquid interface, causing the core to shrink faster than the polymer-free case.

## DISCUSSION

Our work elucidates how bacterial suspensions spatially self-organize into concentric core-shell structures when confined to droplets, driven by the interplay between O_2_ transport, cellular metabolism, and motility. By simultaneously measuring *E. coli* cell concentration, O_2_ concentration, and fluid flows, we found that this self-organization emerges from a feedback mechanism where cellular motility both responds to and shapes O_2_ gradients (Fig. **3**a). By integrating our experiments with theoretical modeling and numerical simulations, we have established a quantitative biophysical framework that describes how this emergent behavior depends on system parameters, such as the droplet size and cell concentration (Fig. **3**i). Our study therefore complements and extends prior work that either mapped O_2_ gradients in non-motile bacterial colonies [29], or explored their consequences in motile suspensions under the influence of gravity, which gives rise to distinct fascinating phenomena [60]. Our work also helps to frame new questions and research directions that build on our approach. For example, owing to the generality of our theoretical framework, it could be extended to other organisms and even mixed systems with multiple cell types and limiting resources [77–82].

### Implications for natural bacterial populations

Aerotaxis is typically thought of as a mechanism by which bacteria can find the most favorable O_2_ conditions for survival and growth [57]. Our work suggests another potential biological function imparted by aerotaxis: it enables cells to reshape their entire population by modulating O_2_ fluxes. For example, we found that under conditions of “transient anoxia” (grey-orange in Fig. **3**i), bacteria use aerotaxis to accumulate at the air-liquid interface and augment O_2_ influx (Fig. **2**c)—enabling cells from the core to eventually recover motility. Exploring the biological implications of this phenomenon, such as for population fitness, will be an interesting direction for future research. Indeed, it could be a previously unrecognized strategy for bacterial populations to optimize their collective resource utilization under confinement. It will also be interesting to explore the biological implications of the core-shell organization revealed by our work more broadly. For example, the formation of an anoxic core could influence other bacterial processes beyond motility—such as quorum sensing [37, 60] and biofilm formation [39]—over long time scales.

Our results could be particularly relevant for understanding bacterial behavior in natural settings, ranging from soils and sediments [83–85] to gels and tissues in the body [5, 17, 37, 64, 86, 87], where cells often experience both spatial confinement and resource limitation [88]. In human health, our results may help shed light on the organization of bacteria in O_2_-limited niches within the body, such as in mucus or chronic wounds, where populations are often layered, as in our experiments. It could be that such hierarchical structures protect a subpopulation of the cells from predators like phages and neutrophils [89] and external stressors such as antibiotics [90, 91]. In environmental science, our findings could help explain bacterial distribution patterns in soil micropores, with potential implications for understanding biogeochemical cycles. The spatial organization we observe could also influence key processes such as carbon mineralization in soil aggregates or nitrogen cycling in marine snow particles, where bacterial activity is often limited by O_2_ availability. Additionally, the principles uncovered by our work could help guide ways to control bacterial populations for practical applications in biotechnology. For example, our findings could be leveraged to design droplet-based bacterial bioreactors or cultivation systems whose spatial organization is controlled to optimize desired metabolic processes (e.g., biofuel synthesis, protein production) while maintaining high cell concentrations and avoiding O_2_ limitations.

### Implications for active matter physics

Our work establishes new principles governing pattern formation in active matter systems—in particular, collectives of actively-moving particles, exemplified in our case by motile bacteria, with spatially varying activity. These are often studied within the context of motility-induced phase separation (MIPS), in which the slowing of particles when they are more concentrated causes them to accumulate, forcing them to spontaneously separate into dilute and dense phases [92]. Our finding that in a large, concentrated droplet, its core acts as a “motility sink” for cells can be thought of as a real-world manifestation of this phenomenon. However, while typical models of MIPS attribute the density-dependence of particle mobility directly to purely mechanical interactions, in our case, changes in particle mobility are coupled to density indirectly, via chemical gradients. Our work demonstrates how this indirect coupling creates a feedback loop where the spatial distribution of activity (bacterial motility) both responds to and shapes the chemical field that controls it—producing new types of non-equilibrium steady states. Our theoretical framework shows how such chemically-mediated feedback can be incorporated into continuum theories of active matter, providing a foundation for studying similar phenomena in other systems.

This study also provides a platform to examine the dynamics of active-passive interfaces (e.g., by monitoring the motile shell-immotile core interface), which has previously been studied only in the context of purely mechanical interactions [93–97]. The additional chemical coupling in our system could introduce additional new effects at these interfaces arising from the interplay between aerotaxis [46], shear-induced depletion [98, 99], and cell-cell hydrodynamic interactions. Such effects might culminate in the emergence of directed long-range flows that can transport nutrients or signaling molecules [100, 101] more efficiently than diffusion [61, 102, 103].

## MATERIALS AND METHODS

For each experiment, we grow an overnight culture of fluorescent *E. coli* (W3110) in LB medium at 30^°^C under shaking conditions. A 1% inoculum is then incubated in fresh LB for 3 h until the optical density (measured using a Biowave Cell Density Meter CO8000) reaches ∼0.5. The cells are washed three times with motility buffer (BMB) and prepared at the desired concentration for a given experiment. The test cell consists of two parallel glass coverslips, cleaned with ethanol before use, separated by a 130 µm-thick air-permeable Parafilm spacer. A droplet of bacterial suspension is deposited on the bottom coverslip, with the volume controlled to reach a desired droplet diameter, with microdroplets of deionized water added around it to maintain humidity and prevent evaporation. The system is sealed with a second overlying coverslip, allowing oxygen to diffuse only through the Parafilm.

We start imaging approximately one minute after the droplet flattens into a disk-like shape, using a Nikon A1R+ inverted laser scanning confocal microscope with a temperature-controlled stage maintained at 30^°^C. Population dynamics are imaged at 1 frame per second using a 4× objective, capturing both bright-field and fluorescence images from an optical slice of ∼100µm thickness. For the state diagram in Fig. **3**i, multiple droplets are imaged using a ∼ 4× objective, with large-area scanning via stitched frames. Flow patterns are visualized using a 20× objective at 30 frames per second from a ~ 10µm optical slice. Dissolved oxygen concentration is measured using a ruthenium complex (Ru(dpp)) whose fluorescence (emission ∼615 nm) is quenched upon oxygen binding. To mitigate toxicity to *E. coli*, the dye is encapsulated in DMPC–PEG2000 phospholipid micelles. Preparation and calibration of the dye follow the protocol by Douarche *et al*. [58].

We use the Beer-Lambert law to calibrate cell concentration in the bright-field images, use particle image velocimetry (PIV) in PIVLab to measure the velocity field generated by the swimming cells, and track individual cells using the ImageJ plugin TrackMate. Due to the non-circular shape of some droplets, we calculated the average intensity profile within a narrow sector of the droplet to obtain the kymographs of cell concentration.

## Acknowledgments

A.M.-C. acknowledges support from the Princeton Center for Theoretical Science, the Princeton Center for the Physics of Biological Function, and the Human Frontier Science Program through the grant LT000035/2021-C. S.S.D. acknowledges support from NSF grants CBET-1941716, DMR-2011750, and EF-2124863 as well as the Camille Dreyfus Teacher-Scholar and Pew Biomedical Scholars Programs, the Eric and Wendy Schmidt Transformative Technology Fund, and the Princeton Catalysis Initiative. We thank Henry Mattingly for assistance with the preparation of the oxygen probe dye; Daniel Amchin, Enkeleida Lushi, and Ned Wingreen for thoughtful discussions; and Daniel Amchin, Rhea Braun, and Nadine Ziegler for assistance with a preliminary version of the experiments at the inception of this project.

## Supporting information (SI)

### Non-dimensionalization of the model

To identify the key governing parameters of our model, we make Eqs. (1a)-(1c) in Fig. **3**a dimensionless. To this end, we choose the O_2_ penetration length 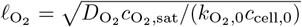, which arises from a balance between O_2_ diffusion and cell O_2_ uptake, as the characteristic length scale. As characteristic O_2_ concentration and cell density we choose the O_2_ saturation concentration, 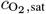, and the initial cell number concentration, *c*_cell,0_, respectively. As characteristic time scale, we consider the time it takes for cells to migrate via aerotaxis over the O_2_ penetration length scale, i.e., 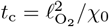. Introducing these characteristic scales, the dimensionless conservation equations for bacteria and O_2_ read:

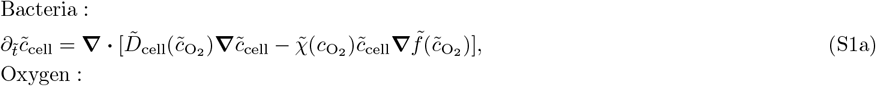

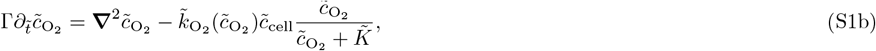

where 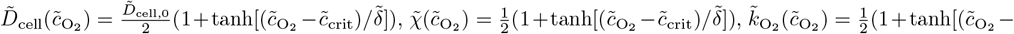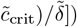, and 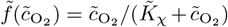. We impose radial symmetry at 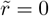, no flux of bacteria across the liquid-air interface at 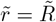, and the dimensionless form of the O_2_ interfacial transport equation (Eq. 1b in Fig. **3**a), given by:

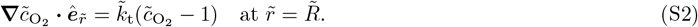

The dimensionless parameters that emerge in Eqs. (S1) and (S2) are:

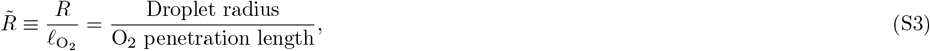

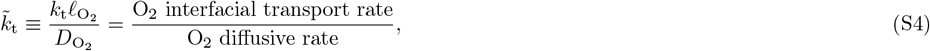

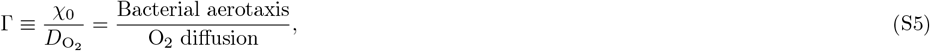

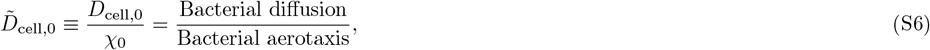

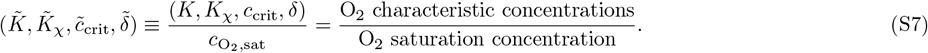

The values of the dimensionless parameters used in the numerical simulations are reported in Table S2. The parameter 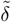, which measures the sharpness of the motility loss transition around the dimensionless critical O_2_ concentration, 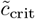, is always set to 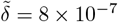.

For the same bacterial species and liquid medium, the only dimensionless parameters that vary in experiments are 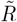 and 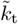. In particular, the parameter 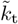, which only enters in the model through the liquid-air boundary condition Eq. (S2), exhibits interesting limiting cases. (i) When 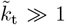, the dimensionless O_2_ concentration at the interface remains constant, i.e., 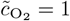, which implies 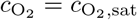. In this limiting case, the interfacial influx of O_2_ is faster than O_2_ diffusion and bacterial O_2_ uptake, so that 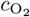 always equilibrates to its saturation value (see Movie S12). However, the influx of O_2_ across the interface is not zero—it adjusts over time to maintain the saturation level of O_2_ at all times. This corresponds to the limit approaching the left in the state diagram in Fig. **3**. (ii) When 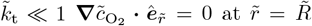, meaning that the influx of O_2_ at the interface is significantly slower than bacterial O_2_ uptake, effectively imposing a no-flux boundary condition (see Movie S13). This limit corresponds to, for example, a scenario with a high density of bacteria that consume O_2_ very efficiently, such that interfacial O_2_ transport cannot keep up. In this case, permanent anoxia should always develop as O_2_ eventually depletes. This corresponds to the limit approaching the right in the state diagram in Fig. **3**.

### Supplementary movies

All movies are available at 10.5281/zenodo.14894704.

**Movie S1**. Population dynamics for a small droplet containing a dilute suspension of bacteria. (Corresponds to main text Fig. 1b.)

**Movie S2**. Population dynamics for a large droplet containing a concentrated suspension of bacteria. (Corresponds to main text Fig. 1d.)

**Movie S3**. High magnification movie of the bacterial motility dynamics within the core and the annulus for a small droplet containing a concentrated bacterial suspension. (Corresponds to main text Fig. 1d magnified panel.)

**Movie S4**. Population dynamics for a small droplet containing a concentrated suspension of bacteria. (Corresponds to main text Fig. 1f.)

**Movie S5**. Simultaneous visualization of the bacterial dynamics (darker means higher cell concentration) and the oxygen concentration field (darker means higher oxygen concentration). The movie is associated with a small concentrated droplet. (Corresponds to main text Fig. 2d.)

**Movie S6**. Bacterial accumulation at the air-liquid interface due to aerotaxis. (Corresponds to main text Fig. 2c.)

**Movie S7**. Population dynamics for a small droplet containing a dilute suspension of bacteria obtained by simulations. (Corresponds to main text Fig. 3b.)

**Movie S8**. Population dynamics for a large droplet containing a concentrated suspension of bacteria obtained by simulations. (Corresponds to main text Fig. 3d.)

**Movie S9**. Population dynamics for a small droplet containing a concentrated suspension of bacteria obtained by simulations. (Corresponds to main text Fig. 3f.)

**Movie S10**. Population dynamics for a small droplet containing a concentrated suspension of bacteria obtained by simulations. Here, we have increased the diffusion coefficient and the aerotactic sensitivity of bacteria by a factor of 4 which is comparable to values in the viscoelastic fluid case.

**Movie S11**. Population dynamics for a small viscoelastic droplet containing a concentrated suspension of bacteria. The continuous phase is an aqueous polymeric solution PEO 5 MDa *c*_pol_ = 0.2wt.%. (Corresponds to main text Fig. 4.)

**Movie S12**. Left: Dimensionless bacterial and O_2_ concentrations as a function of the dimensionless radius over time, obtained by simulations. Right: Dimensionless O_2_ concentration, O_2_ influx, and cell concentration at the droplet interface as functions of dimensionless time. Here, 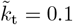 and 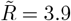.

**Movie S13**. Same as Movie S12 but for 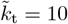 and 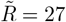.

---

**Fig. S1:**
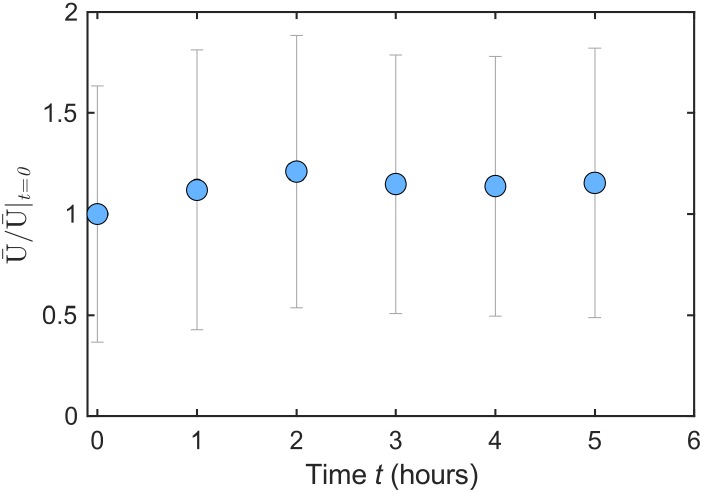
Mean speed of bacteria measured in BMB over 5 hours. We track individual bacteria in a dilute suspension with *c*_cell,0_ = 10^8^ cells*/*mL every hour, over a period of five hours. As shown by the data, the mean swimming speed BMB does not change, indicating that the cells have enough endogenous energy resources to continue swimming over this duration, consistent with prior measurements [1]. The error bars correspond to the standard deviation of the measured speed distribution.

**Fig. S2:**
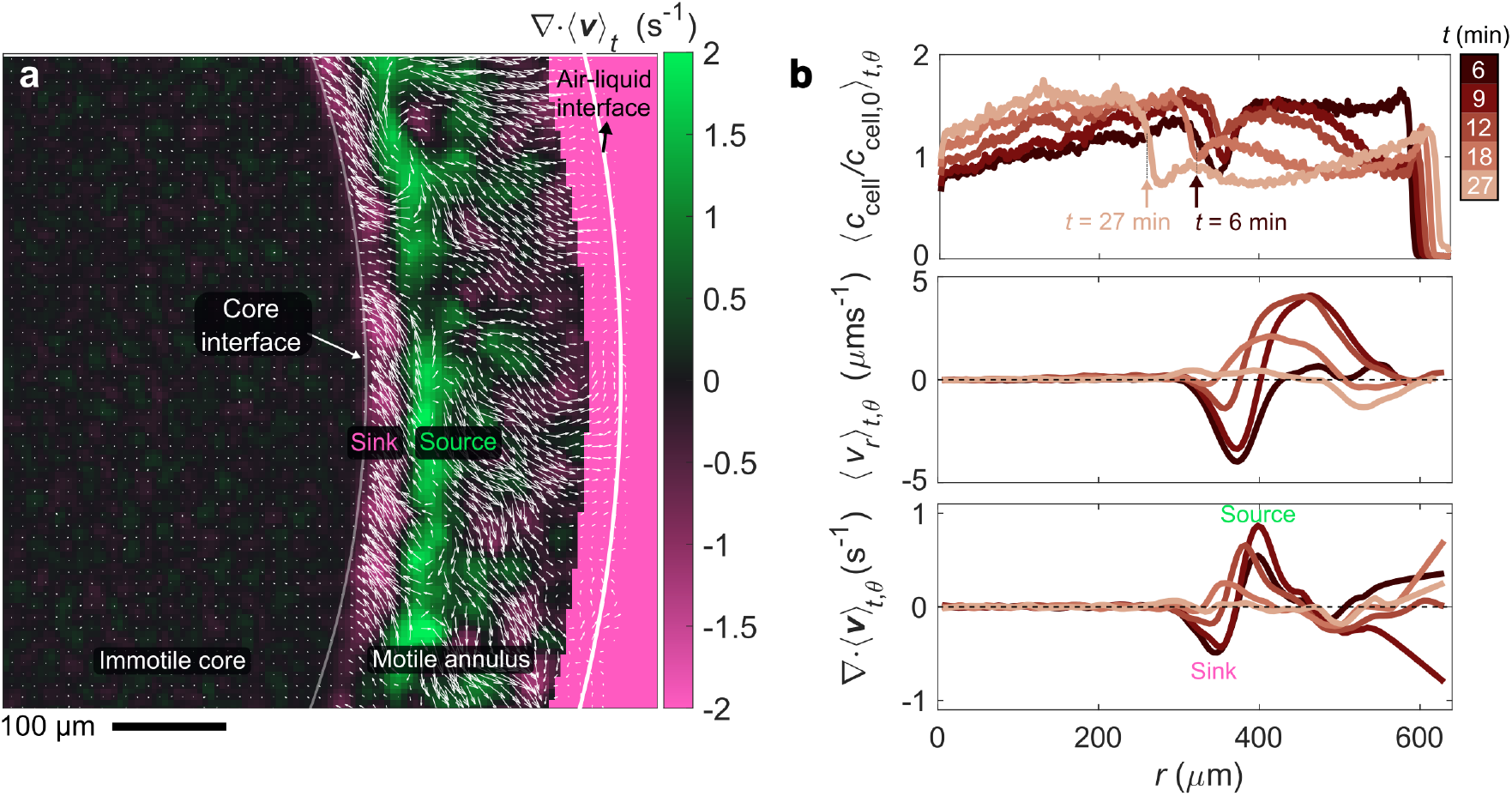
Mean flow field generated by swimming bacteria in the annulus. **a**, Mean flow field generated for *c*_cell,0_ = 8 × 10^10^cells*/*mL measured at a height of 30 µm from the bottom surface. The background colormap shows the divergence of the mean velocity field **∇** · ⟨***v***⟩ _*t,θ*_. **b** Time evolution of the normalized cell concentration ⟨*c*_cell_*/c*_cell,0_⟩ _*t,θ*_ (top), the mean radial component of the velocity *v*_*r t,θ*_ (middle), and the divergence of the mean velocity field ∇ · ⟨***v***⟩_*t,θ*_ (bottom). The mean velocity field and all the mean profiles are obtained by averaging over 1 minute (1800 frames). The arrows in the top panel show the location of the core boundary at *t* = 6 and 27 minutes after the start of the experiment. The divergence of the velocity field shows a positive value at a radial distance 100 µm away from the core boundary. This observation indicates that there is a vertical top-down flow, acting as a *source* of cells, that splits into two radial streams close to the bottom surface. The stream between the *source* and the core is directed towards the core (⟨*v*_*r*_⟩_*t*_ *<* 0) where they lose their motility after *τ*_delay_ ≈ 5 minutes [2], sediment and locally increase the *c*_cell_. In this part of the annulus, we observe ∇ · ⟨***v***⟩_*t*_ *<* 0, *i*.*e*., the core is acting as another *sink* for cells. The opposite stream moves in the radial direction towards the air-liquid interface (⟨*v*_*r*_ ⟩_*t*_ *>* 0) resulting in an accumulation at the interface again acting as a *sink* for the flux of bacteria. Thus, with sinks pulling cells both towards the core and the air-liquid interface, the middle part of the annulus gradually becomes depleted of bacteria.

**TABLE S1:** Diffusion coefficient of individual cells at different PEO concentrations. We first prepare an overnight culture of bacteria. Next, these bacteria are re-suspended at a more dilute concentration of 3 to 5 × 10^5^ cells/mL in the polymer solution of choice. Next, the suspension of cells is put between two glass slides separated by paraffin film. The cells are then imaged with the confocal microscope with a 10 × air objective at a temperature of 30^°^C. We acquire 512 by 512 pixel images at ≈33 ms time intervals. We use the ImageJ plugin TrackMate to track bacteria and we write a custom MATLAB script to extract speed and running time distributions. From these distributions we compute the average bacterial speed, 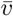, and average run duration 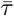. We then estimate the diffusion coefficient as 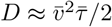 following [3].

| $c$ (w/w%) | $\bar{v}$ (μm/sec) | $\bar{\tau}$ (sec) | $D$ (μm<sup>2</sup>/sec) |
| --- | --- | --- | --- |
| 0 | 25 | 0.6 | 187 |
| 0.1 | 40 | 0.6 | 480 |
| 0.15 | 38 | 0.6 | 433 |
| 0.2 | 43 | 0.8 | 725 |

**Fig. S3:**
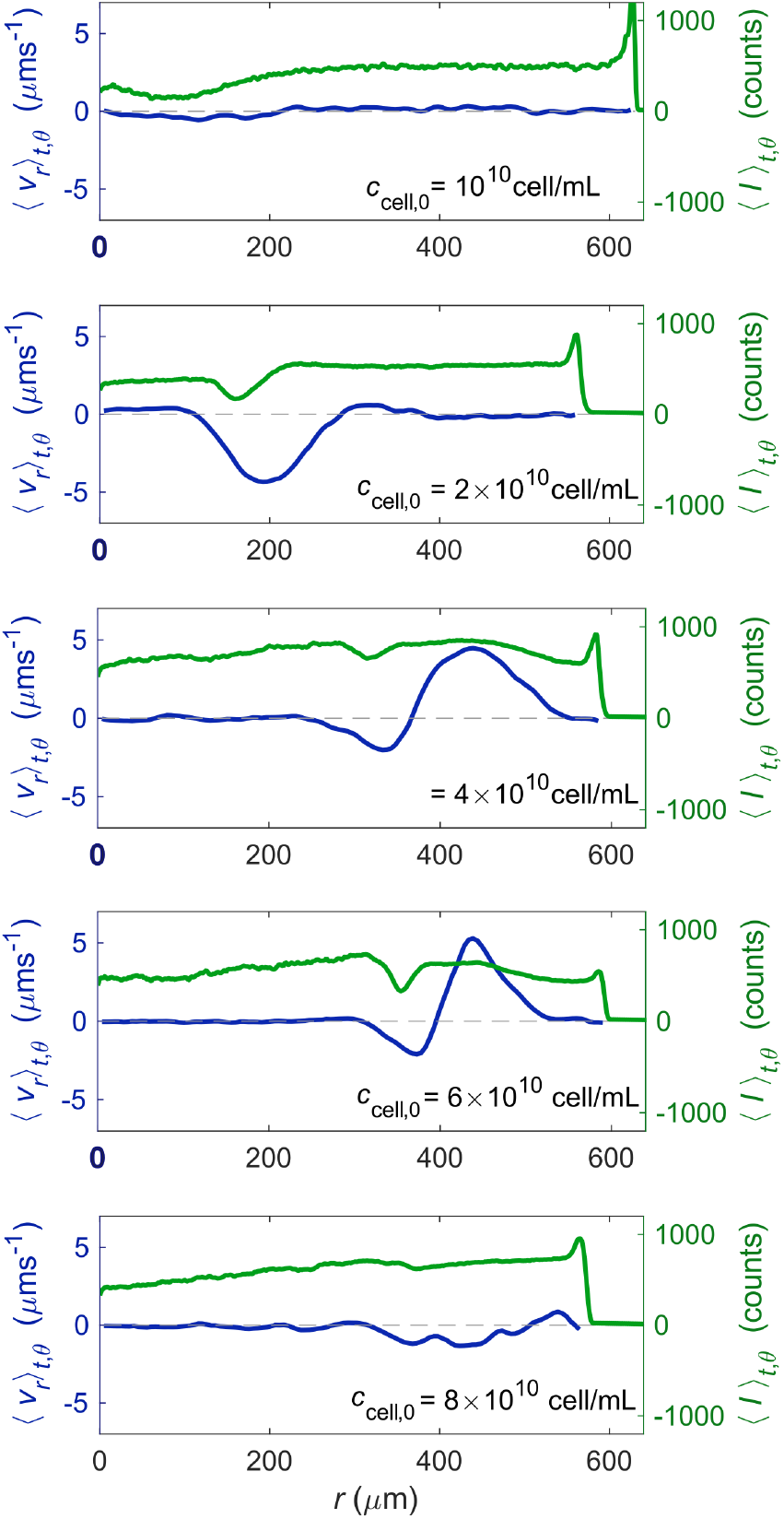
Effect of cell concentration on the mean flow field generated by bacteria. The left axis shows the radial component of the velocity field plotted against the radial direction. On the right axis, the instantaneous GFP signal intensity profiles are displayed in light grey, with the mean intensity profile highlighted in orange. Since intensity is proportional to cell concentration, the small dip in the intensity profile indicates the edge of the core. The mean profiles are calculated by averaging data recorded during the first two minutes following core formation.

**Fig. S4:**
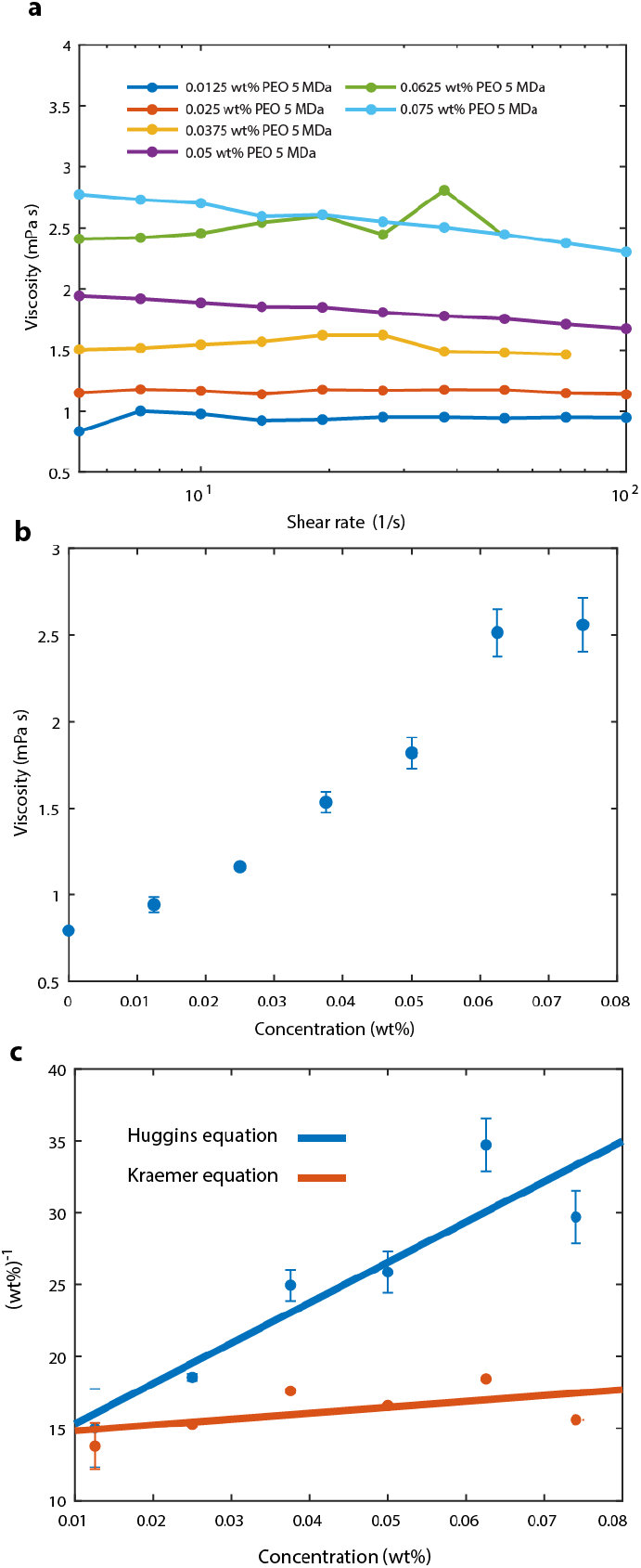
Rheological characterization of polymer solutions. We use an Anton Paar MCR501 rheometer with a double-gap Couette geometry to characterize our polymer solutions. **a** We measure the dynamic shear viscosity as a function of imposed shear rate; as shown by the data, for the polymer concentrations used in this study we observe shear-thinning behavior. **b** Low shear rate viscosity vs. concentration curve. The dots represent the mean viscosity measured over all shear rates at constant polymer concentration, while the error bars correspond to the variation of the viscosity at constant polymer concentration over the range of shear rates used, 5 − 200 1/s. This dependence of viscosity on polymer concentration *c* yields a straightforward way to determine the polymer overlap concentration *c*^*^. In particular, writing the viscosity as a virial expansion 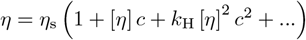 and retaining only the first three terms on the right hand side yields two distinct relations [4]; here, [*η*] is known as the intrinsic viscosity, *k*_H_ is a constant known as the Huggins coefficient, and *η*_s_ is the viscosity of the solvent. The first, known as the Huggins equation, directly follows from rearranging this equation: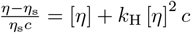. The second, known as the Kraemer equation, follows from the approximation that ln 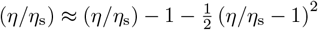; substituting from the virial expansion for *η/η*_s_ then yields the Kraemer equation: 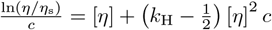. **c** Fitting both Huggins and Kraemer equations to our data and extrapolating to *c* = 0 then yields a direct determination of the intrinsic viscosity; we then use the established relation *c*^*^ = 0.77*/* [*η*] [5] to directly compute *c*^*^, which renders a value of 0.05 ± 0.03 w/w% for PEO 5 MDa dissolved in BMB.

**TABLE S2:**
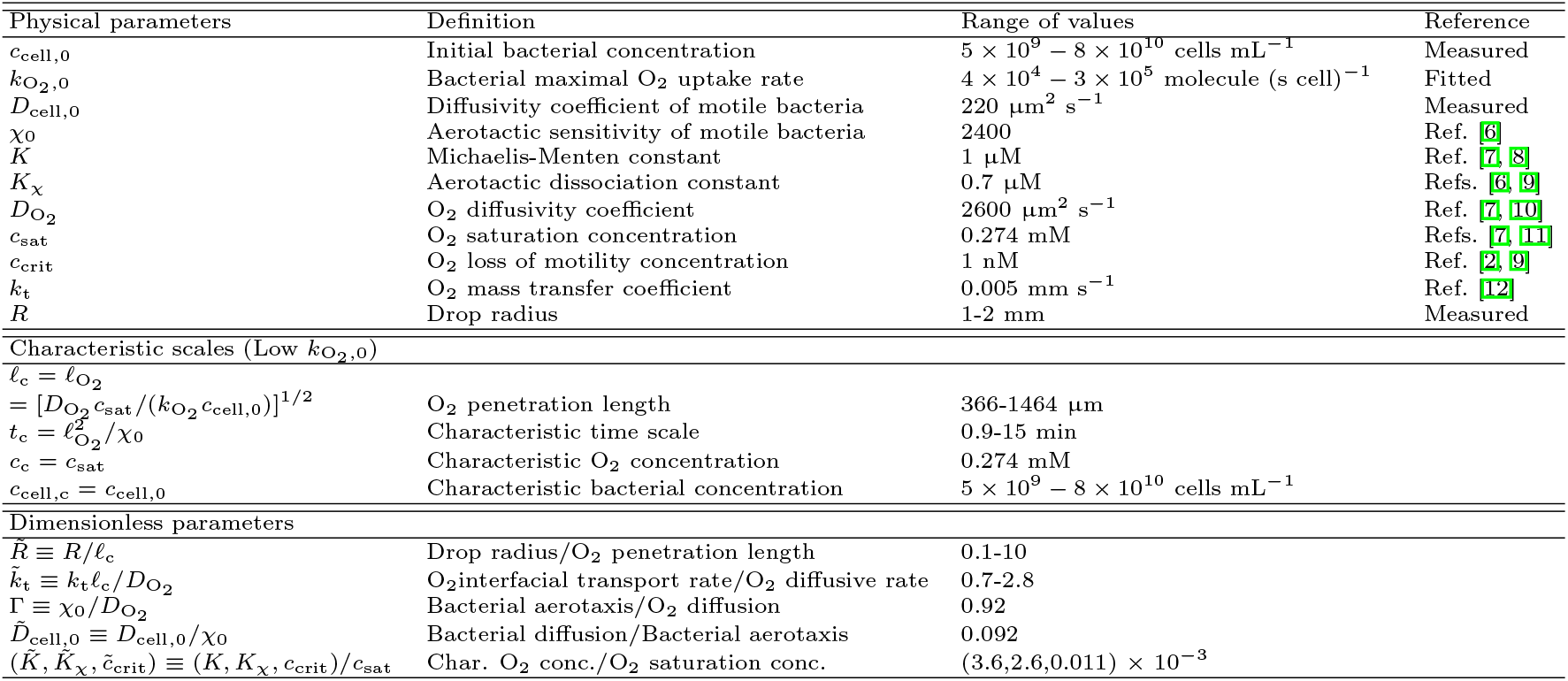
Estimates and experimental measurements of the physical and dimensionless parameters used in the model.

